# New β-Lactamase Inhibitors Nacubactam and Zidebactam Improve the In Vitro Activity of β-Lactam Antibiotics Against *Mycobacterium abscessus* Complex Clinical Isolates

**DOI:** 10.1101/597765

**Authors:** Amit Kaushik, Nicole C. Ammerman, Nicole M. Parrish, Eric L. Nuermberger

## Abstract

The new diazabicyclooctane-based β-lactamase inhibitors avibactam and relebactam improve the in vitro activity of β-lactam antibiotics against *Mycobacterium abscessus* complex (MABC). Here, we evaluated the in vitro activity of two newer diazabicyclooctane-based β-lactamase inhibitors in clinical development, nacubactam and zidebactam, with β-lactams against clinical isolates of MABC. Both inhibitors lowered the MICs of their partner β-lactams, meropenem (eight-fold) and cefepime (two-fold), and those of other β-lactams, similar to prior results with avibactam and relebactam.

## Introduction

*Mycobacterium abscessus* complex (MABC) is comprised of rapidly growing, nontuberculous mycobacteria responsible for chronic, difficult-to-treat lung, skin, and wound infections that are increasing in prevalence (1–4). Both intrinsic and acquired drug resistance contribute to the recalcitrance of MABC lung infections (5). Despite the outstanding contribution of β-lactam antibiotics to treatment of infectious diseases, their utility against MABC organisms is limited by a chromosomally encoded, broad-spectrum, Ambler class A β-lactamase, BlaMab, which is the major determinant of intrinsic β-lactam resistance in MABC (6). While older β-lactam-based β-lactamase inhibitors (BLIs) such as clavulanate, tazobactam and sulbactam are ineffective against BlaMab and do not improve the in vitro activity of β-lactam antibiotics against MABC organisms (7, 8), we and others have shown that the new diazabicyclooctane-based BLIs avibactam and relebactam, developed to treat multidrug-resistant Gram-negative bacteria (9), do improve the in vitro activity of many β-lactam antibiotics against MABC organisms, particularly carbapenems and cephalosporins (8, 10–12). Avibactam and relebactam have been developed with ceftazidime and imipenem, respectively. However, ceftazidime has poor intrinsic activity against MABC organisms, as evidenced by high MICs despite combination with avibactam or relebactam (10, 12), while imipenem has relatively high intrinsic activity and MICs are only modestly lower in the presence of these BLIs (8, 10). Newer diazabicyclooctane-based BLIs being developed for treatment of challenging Gram-negative infections, including nacubactam and zidebactam (13, 14), may offer advantages over avibactam and relebactam. Both nacubactam (OP0595, RG6080) co-formulated with meropenem and zidebactam (WCK 5107) co-formulated with cefepime (co-formulation is WCK 5222) have completed clinical safety, tolerability, pharmacokinetics and lung penetration studies (ClinicalTrials.gov identifiers: NCT02972255, NCT03182504, NCT02674347, NCT03630094) and received Fast Track and Qualified Infectious Disease Product (QIDP) designations from the U.S. Food and Drug Administration (15, 16). The aim of our study was to evaluate the activity of nacubactam or zidebactam in combination with β-lactams against drug-resistant clinical isolates of MABC.

## Materials and Methods

Nacubactam and zidebactam were procured from MedKoo Biosciences, Inc., NC, USA (purity >98%). A total of twenty-six β-lactam antibiotics (Table 1), including penicillins, cephalosporins and carbapenems, were purchased from commercial sources as previously described (10). The purity of all β-lactams was >95%. All drugs were stored and dissolved either in DMSO or water prior to drug susceptibility testing (DST) according to manufacturers’ recommendation.

**TABLE 1.** MIC values of β-lactams with and those without β-lactamase inhibitors against *M. abscessus* subsp. abscessus strain ATCC 19977 in Middlebrook 7H9 medium

| | MIC in $\mu\text{g/mL}$ | | | | |
| --- | --- | --- | --- | --- | --- |
|  | Alone | With nacubactam |  | With zidebactam |  |
| β-lactam tested | | 4 $\mu\text{g/mL}$ | 8 $\mu\text{g/mL}$ | 4 $\mu\text{g/mL}$ | 8 $\mu\text{g/mL}$ |
| **Oral carbapenems** |  |  |  |  |  |
| Faropenem | 128 | 32 | 32 | 32 | 32 |
| Tebipenem | 256 | 8 | 4 | 16 | 16 |
| **Parenteral carbapenems** |  |  |  |  |  |
| Biapenem | 16 | 4 | 4 | 4 | 4 |
| Doripenem | 16 | 4 | 2 | 4 | 4 |
| Ertapenem | >256 | 16 | 16 | 64 | 64 |
| Imipenem | 8 | 4 | 2 | 2 | 2 |
| Meropenem | 16 | 4 | 2 | 8 | 8 |
| **Oral cephalosporins** |  |  |  |  |  |
| Cefdinir | 32 | 16 | 16 | 16 | 16 |
| Cefixime | >256 | 128 | 128 | 256 | 128 |
| Cefpodoxime | >256 | 64 | 64 | 128 | 64 |
| Cefuroxime<sup>a</sup> | 128 | 8 | 8 | 16 | 16 |
| Cephalexin | >256 | >256 | >256 | >256 | >256 |
| **Parenteral cephalosporins** |  |  |  |  |  |
| Cefazolin | >256 | >256 | 256 | >256 | >256 |
| Cefepime | 32 | 32 | 16 | 16 | 16 |
| Cefoperazone | >256 | >256 | >256 | >256 | >256 |
| Cefotaxime | 128 | 64 | 32 | 64 | 64 |
| Cefoxitin | 32 | 32 | 32 | 32 | 32 |
| Ceftaroline | >256 | 8 | 8 | 64 | 32 |
| Ceftazidime | >256 | >256 | >256 | >256 | >256 |
| Ceftriaxone | >256 | 32 | 16 | 128 | 32 |
| Cephalothin | >256 | 256 | 128 | >256 | >256 |
| Moxalactam | 128 | 128 | 128 | 128 | 128 |
| **Penicillins** |  |  |  |  |  |
| Amoxicillin | >256 | 16 | 16 | 256 | 256 |
| Cloxacillin | >256 | >256 | >256 | >256 | >256 |
| Dicloxacillin | >256 | >256 | >256 | >256 | >256 |
| Oxacillin | >256 | >256 | >256 | >256 | >256 |

270 <sup>a</sup>Cefuroxime is available in both oral and parenteral formulations.

Twenty-eight clinical isolates of MABC were collected at Johns Hopkins Hospital, Baltimore, MD, USA from 2005 to 2015 and described previously (8, 10). *M. abscessus* ATCC 19977 was purchased from the American Type Culture Collection (Manassas, VA, USA) and used as a reference strain. Middlebrook 7H9 broth supplemented with 10% Middlebrook OADC enrichment, 0.5% glycerol, and 0.05% Tween 80, was used as the growth medium. Middlebrook 7H9 broth supplemented with 10% OADC and 0.5% glycerol was used primarily for minimum inhibitory concentration (MIC) determination instead of cation-adjusted Mueller-Hinton broth (CAMHB) because growth of clinical isolates is faster in Middlebrook 7H9 broth compared to CAMHB, thus limiting the potential for over-estimation of MICs due to β-lactam instability in the medium, as discussed previously (10).

MIC was determined using the microbroth dilution method in round bottom wells in 96-well plates, as previously described (8, 10). In brief, 100 μL of media was dispensed in wells. Drugs were dissolved and two-fold dilutions were prepared ranging from 2 to 256 μg/mL. Wells were prepared with β-lactams alone or in combination with a fixed concentration of 4 or 8 μg/mL of either nacubactam or zidebactam, or either BLI alone. A total of 100 μL of a log phase culture containing 1 × 10^4^ to 5 × 10^4^ CFU was added to each well except the negative control well (media only). Plates were incubated at 30°C for 3 days for Middlebrook 7H9 broth. The MIC was defined as the lowest concentration of β-lactam that prevented growth as observed by the naked eye. MIC_50_ and MIC_90_ were defined as the MIC at which at least 50% and at least 90%, respectively, of the clinical MABC isolates were inhibited. DST was repeated to confirm the MIC against *M. abscessus* ATCC 19977.

## Results

Initially, we studied the effect of β-lactams in presence and absence of nacubactam and zidebactam against *M. abscessus* ATCC 19977. Both BLIs improved the activity of carbapenems and some cephalosporins (Table 1). The potentiating effects were greatest with tebipenem, ertapenem, cefuroxime, ceftaroline and, to a lesser extent, meropenem. However, nacubactam was generally slightly more effective than zidebactam and it uniquely potentiated the effects of amoxicillin. Nacubactam at 8 μg/mL resulted in two-fold lower MICs compared to 4 μg/mL for some β-lactams, while zidebactam results were similar irrespective of the concentration tested. Specifically, nacubactam at 8 μg/mL and zidebactam at 4-8 μg/mL improved the activity of their partner β-lactams, meropenem and cefepime by eight-fold and two-fold, respectively. As previously observed with avibactam and relebactam, MICs of cefoxitin remained unchanged in the presence of nacubactam and zidebactam, reflecting the stability of cefoxitin to MABC β-lactamase activity (17). The MICs of nacubactam and zidebactam against *M. abscessus* 19977 was >256 μg/mL, suggesting that their potentiation of β-lactam activity were due to β-lactamase inhibition rather than any intrinsic anti-bacterial effects.

We chose 8 μg/mL for nacubactam and 4 μg/mL for zidebactam as fixed concentrations to screen against the clinical isolates. On average, the clinical isolates were more resistant than *M. abscessus* 19977. However, both BLIs improved the activity of selected β-lactams (Table 2, Figures 1 and 2). Nacubactam and zidebactam lowered the MIC_50_ values of their partner β-lactams, meropenem and cefepime by 8-fold and 2-fold, respectively, as well as those of the carbapenems, several cephalosporins (ceftaroline, cefuroxime and cefdinir) and, in the case of nacubactam, amoxicillin, consistent with their effects against ATCC 19977.

**Table 2.** MIC values of β-lactams with and those without nacubactam or zidebactam against 28 drug-resistant MABC clinical isolates in Middlebrook 7H9 medium

| MICs ($\mu\text{g/mL}$) | | | | | | | | | |
| --- | --- | --- | --- | --- | --- | --- | --- | --- | --- |
|  | Alone |  |  | With nacubactam<sup>a</sup> |  |  | With zidebactam<sup>a</sup> |  |  |
|  | Range | MIC<sub>50</sub> | MIC<sub>90</sub> | Range | MIC<sub>50</sub> | MIC<sub>90</sub> | Range | MIC<sub>50</sub> | MIC<sub>90</sub> |
| **Oral carbapenem** |  |  |  |  |  |  |  |  |  |
| Tebipenem | 64 - >256 | 256 | >256 | 4 - 32 | 8 | 16 | 16 - 256 | 32 | 128 |
| **Parenteral carbapenems** |  |  |  |  |  |  |  |  |  |
| Biapenem | 8 - 256 | 16 | 64 | 4 - 8 | 8 | 8 | 4 - 64 | 8 | 32 |
| Doripenem | 8 - 128 | 32 | 64 | 4 - 16 | 8 | 8 | 4 - 64 | 4 | 32 |
| Ertapenem | 128 - >256 | 256 | >256 | 8 - 64 | 16 | 64 | 16 - >256 | 64 | 256 |
| Imipenem | 8 - 64 | 16 | 32 | 4 - 16 | 8 | 16 | 4 - 32 | 8 | 16 |
| Meropenem | 8 - 256 | 32 | 256 | 4 - 16 | 4 | 8 | 4 - 128 | 8 | 64 |
| **Oral cephalosporins** |  |  |  |  |  |  |  |  |  |
| Cefdinir | 32 - 256 | 64 | 128 | 16 - 32 | 16 | 32 | 16 - 64 | 32 | 64 |
| Cefuroxime<sup>b</sup> | 64 - >256 | 256 | >256 | 8 - 32 | 16 | 32 | 16 - 256 | 32 | 64 |
| **Parenteral cephalosporins** |  |  |  |  |  |  |  |  |  |
| Cefepime | 16 - 128 | 32 | 64 | 8 - 64 | 16 | 32 | 8 - 64 | 16 | 32 |
| Cefoxitin | 32 - 64 | 32 | 64 | 32 - 64 | 32 | 64 | 32 - 64 | 32 | 64 |
| Ceftaroline | 64 - >256 | >256 | >256 | 4 - 32 | 8 | 16 | 16 - >256 | 64 | 256 |
| **Oral penicillin** |  |  |  |  |  |  |  |  |  |
| Amoxicillin | >256 - >256 | >256 | >256 | 8 - 256 | 16 | 64 | 64 - >256 | 256 | >256 |

<sup>a</sup>Nacubactam and zidebactam were used at fixed concentrations of 8 and 4 $\mu\text{g/mL}$ , respectively.
<sup>b</sup>Cefuroxime is available in both oral and parenteral formulations.

**Figure 1.**
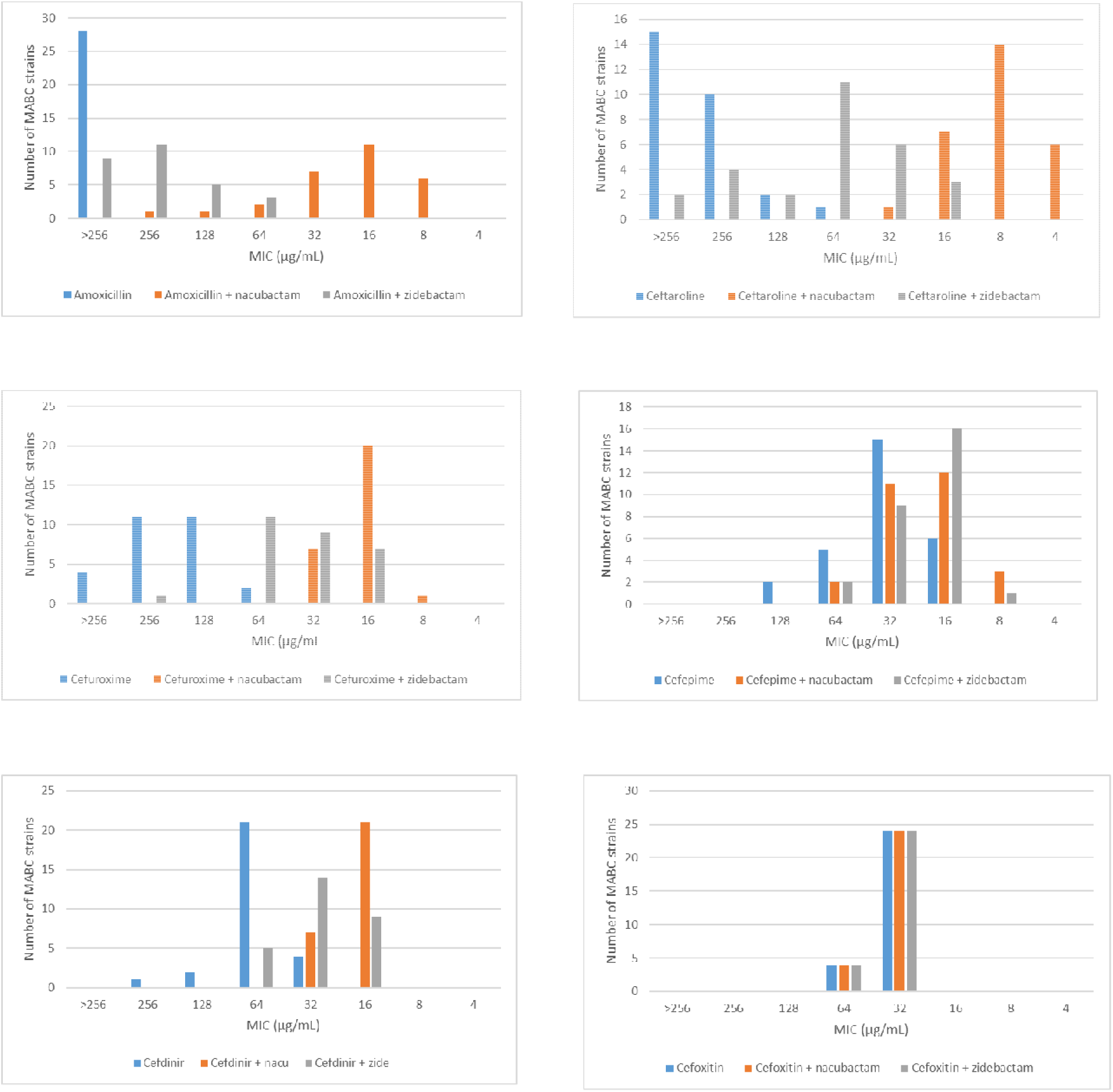
MIC distributions of amoxicillin and cephalosporins, alone and in combination with 8 μg/ml nacubactam or 4 μg/ml zidebactam, against 28 MABC clinical isolates.

**Figure 2.**
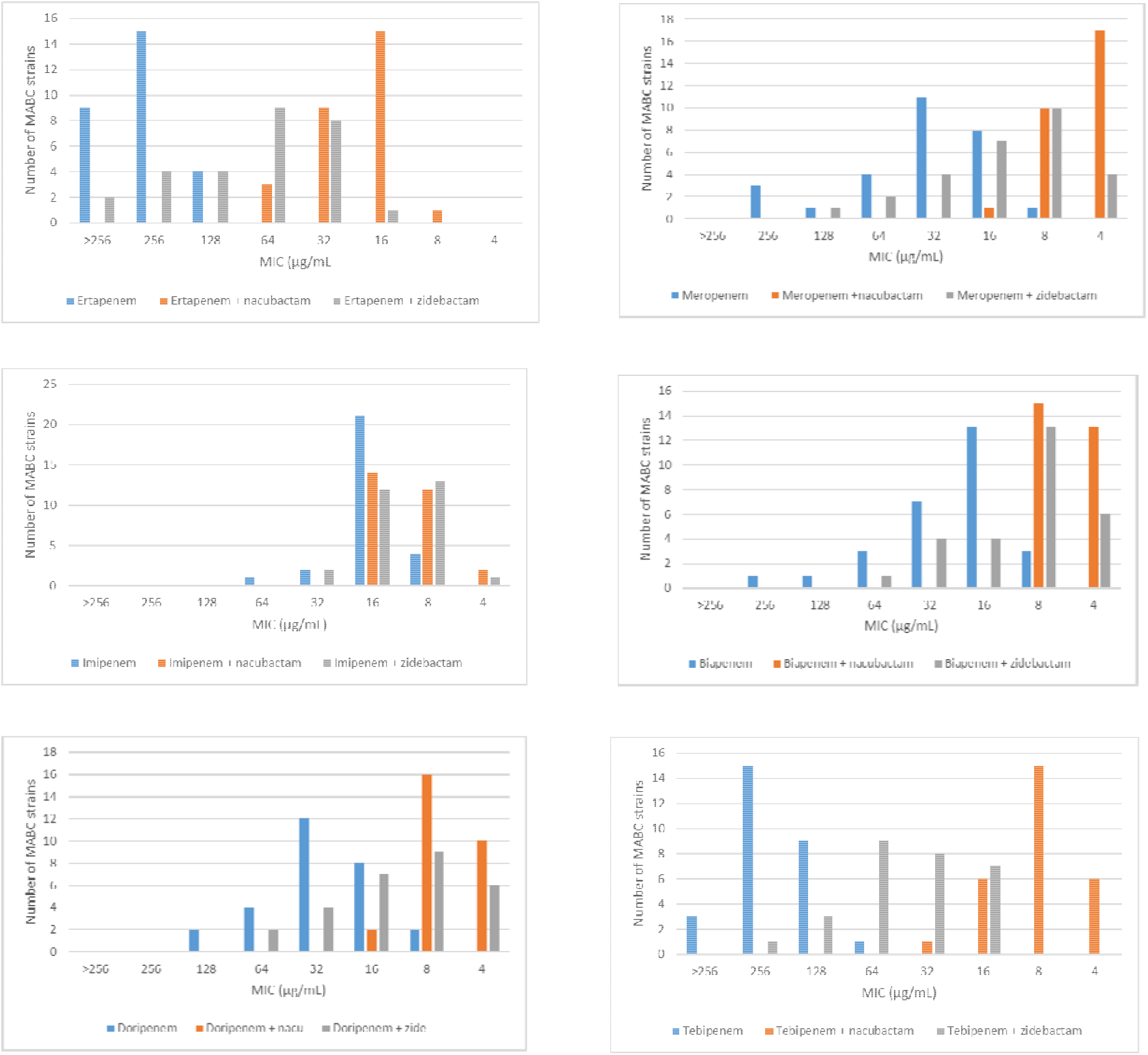
2 MIC distributions of carbapenems, alone and in combination with 8 μg/ml nacubactam or 4 μg/ml zidebactam, against 28 MABC clinical isolates.

Against the clinical isolates, the addition of 8 μg/mL nacubactam reduced the meropenem MIC_50_ from 32 μg/mL to 4 μg/mL, thus changing the interpretation from resistant to susceptible, according to CLSI breakpoints for *M. abscessus* (albeit using 7H9 broth rather than the CAMHB media recommended by CLSI, for reasons we explained previously) (10). Indeed, all 28 clinical isolates had MICs within the susceptible-to-intermediate range when meropenem was combined with nacubactam. These results are somewhat better than those observed in our previous study when meropenem was combined with vaborbactam 4 μg/mL (10).

## Discussion

For β-lactams, the percentage of the dosing interval for which free drug concentrations exceed the MIC μg/mL (%fT_>MIC_) is the pharmacokinetic/pharmacodynamic parameter best correlated with antibacterial effect (18). Target values for %fT_>MIC_ vary among sub-classes of β-lactams and by organism. Although such targets are not established for β-lactams against MABC organisms, target %fT_>MIC_ values against other bacteria are ≈40% for carbapenems and ≈40-60% for cephalosporins (19, 20). Monogue et al showed that nacubactam plasma concentrations exceed 8 μg/mL for about 60% of the dosing interval when dosed intravenously at 1.5 grams every 8 hours (0.5 hr infusion) (13), suggesting that β-lactam MICs in the presence of nacubactam 8 μg/mL may predict clinical efficacy if the β-lactam dosing regimen meets the %fT_>MIC_ target for MIC in the presence of the BLI. Likewise, susceptibility breakpoints based on such targets should be predictive of clinical efficacy. Although no breakpoint has been established for cefepime against MABC organisms, the addition of zidebactam 4 μg/mL (or nacubactam 8 μg/mL) reduced the cefepime MIC_50_ from the resistant to the intermediate susceptibility range when considering the CLSI breakpoints for cefepime against *Pseudomonas aeruginosa* (21, 22). Zidebactam plasma and alveolar epithelial lining fluid concentrations exceed 4 μg/mL for at least 75% and at least 50%, respectively, of the dosing interval when cepepime/zidebactam are dosed intravenously at 2g/1gevery 8 hours (1 hr infusion) in healthy subjects (16).

In conclusion, this study demonstrates that nacubactam and zidebactam improve the anti-MABC activity of carbapenems, several cephalosporins, and, in the case of nacubactam, amoxicillin. Specifically, addition of nacubactam lowered meropenem MICs eight-fold, resulting in all isolates being susceptible or intermediately susceptible by CLSI interpretive criteria for meropenem. In our previous study (10), the meropenem/vaborbactam combination was not quite as potent as the meropenem/nacubactam combination studied here against the same isolates, suggesting that meropenem/nacubactam, if approved, could have an advantage for the treatment of MABC infections. However, further head-to-head comparisons with larger numbers of clinical isolates are required before drawing a more confident conclusion. Zidebactam had a more modest effect on cefepime MICs and cefepime has lower intrinsic activity against MABC than meropenem. However, emerging evidence suggests that combinations of two β-lactams with an effective BLI could be synergistic against *M. abscessus* (12, 23, 24). Our study identified β-lactams belonging to several sub-classes that are potentiated by new BLIs and could be combined with a fixed β-lactam/BLI combination to pursue such synergistic effects.

## Acknowledgements

The authors gratefully acknowledge Dr. Gyanu Lamichhane for providing partial characterization of the MABC clinical isolates. Funding was provided by the National Institutes of Health, R21AI137814 (ELN).

